# Deletion of the Wnt regulator Znrf3 alters bone geometry without inducing high bone mass

**DOI:** 10.64898/2026.03.30.715366

**Authors:** Cassandra R Diegel, Megan N Michalski, Gabrielle Foxa Wiartalla, Zhendong A Zhong, Zachary B Madaj, Bart O Williams

## Abstract

RNF43 and ZNRF3 are transmembrane E3 ubiquitin ligases that negatively regulate Wnt signaling by promoting ubiquitination and degradation of Frizzled receptors. Loss of either gene enhances Wnt/β-catenin signaling and has been linked to tumorigenesis. Wnt signaling is a key regulator of skeletal development and bone homeostasis, and pharmacologic activation of this pathway is an established therapy for osteoporosis. In *Xenopus laevis*, simultaneous disruption of *rnf43* and *znrf3* results in supernumerary limb formation; however, their roles in mammalian limb development and skeletal maintenance remain unclear.

We demonstrate that mice homozygous for null alleles of both *Rnf43* and *Znrf3* do not develop supernumerary limbs. Because activation of Wnt/β-catenin signaling in osteoblasts increases bone mass, we hypothesized that osteoblast-specific deletion of *Rnf43* and/or *Znrf3* would produce a high-bone-mass phenotype. Instead, osteoblast-specific loss of *Znrf3* resulted in age-and sex-dependent reductions in trabecular bone mass, characterized by decreased bone mineral density and bone volume fraction, reduced trabecular number, and increased trabecular separation. Cortical bone exhibited increased cross-sectional size with reduced cortical area fraction and altered structural properties, while tissue mineral density was unchanged. In contrast, deletion of *Rnf43* had minimal skeletal effects, and combined deletion of both genes did not exacerbate the phenotype observed with loss of *Znrf3* alone.

These findings identify *Znrf3* as the dominant functional paralog regulating bone architecture in mature osteoblasts and underscore the importance of evaluating skeletal geometry when modulating upstream Wnt regulators.

## INTRODUCTION

Wnt signaling is a central regulator of skeletal development and adult bone homeostasis^1–3^. Precise modulation of canonical Wnt/β-catenin activity within the osteochondral lineage governs bone mass acquisition, trabecular architecture, and cortical geometry. Genetic activation of Wnt signaling in osteoblasts, through stabilization of β-catenin or deletion of pathway inhibitors such as *Apc,* produces dramatic high-bone-mass phenotypes^4^. Conversely, attenuation of the pathway leads to bone loss^4–6^. These observations have led to the development of Wnt-targeting therapeutics, including the FDA-approved osteoporosis treatment Romosozumab^7–9^, underscoring the clinical relevance of defining how Wnt pathway components regulate skeletal biology.

At the level of the cell surface, Wnt signal strength is tightly controlled by the transmembrane RING-type E3 ubiquitin ligases, Ring Finger Protein 43 (RNF43) and Zinc and Ring Finger Protein 3 (ZNRF3). These proteins promote ubiquitination and internalization of Frizzled (FZD) receptors, thereby limiting cellular responsiveness to Wnt ligands^10, 11^. Their activity is antagonized by R-spondin (RSPO) proteins, which bind RNF43/ZNRF3 in conjunction with LGR4/5/6 receptors and induce clearance of the complex from the plasma membrane, increasing FZD abundance and potentiating Wnt signaling^10^. Dysregulation of the RSPO–RNF43/ZNRF3 module has been implicated in cancer^12^ and developmental disorders^10^, highlighting its importance in tissue homeostasis.

Although RNF43 and ZNRF3 are often described as functionally redundant, genetic studies suggest context-dependent roles. *Rnf43*-deficient mice are viable and lack overt developmental abnormalities^11^, whereas *Znrf3-*deficient mice die perinatally with defects including abnormal lens development and partially penetrant defects in neural tube closure^13^. In Xenopus, simultaneous disruption of *rnf43* and *znrf3* results in supernumerary limb formation, implicating these ligases in early limb patterning^14^. However, the consequences of combined loss of *Rnf43* and *Znrf3* in mammals, particularly within the skeletal system, have not been fully defined. Given the established importance of Wnt signaling in bone, determining how receptor-level regulation by RNF43 and ZNRF3 influences skeletal development and homeostasis remains an important unresolved question.

In this study, we investigated the roles of *Rnf43* and *Znrf3* in murine skeletal biology using both germline and osteoblast-specific conditional knockout models. We first asked whether combined germline deletion recapitulates the supernumerary limb phenotype observed in amphibians. We then examined whether deletion of these negative Wnt regulators in mature osteoblasts would enhance Wnt signaling sufficiently to produce a high-bone-mass phenotype, as predicted from canonical Wnt gain-of-function models. Unexpectedly, combined germline deletion of *Rnf43* and *Znrf3* in mice did not induce limb duplication. Moreover, osteoblast-specific deletion of *Znrf3*, but not *Rnf43*, resulted in age- and sex-dependent reductions in trabecular bone mass and alterations in cortical geometry without changes in tissue mineral density. Simultaneous deletion of both genes in mature osteoblasts did not exacerbate the phenotype observed with loss of *Znrf3* alone. These findings identify *Znrf3* as the dominant functional paralog regulating skeletal architecture in mature osteoblasts and demonstrate that receptor-level modulation of Wnt signaling produces outcomes distinct from downstream β-catenin stabilization.

## RESULTS

### Loss of *Rnf43* and *Znrf3* in the Germline Does Not Induce Supernumerary Limb Formation

To generate mice carrying global (germline) deletions of *Rnf43* and *Znrf3*, mice harboring floxed alleles (*Rnf43^Flox^* and *Znrf3^Flox^*)^11^ were crossed to CMV-Cre^15^. After confirming Cre-mediated recombination in the germline, we characterized individual knockout lines. As previously reported, *Rnf43^KO/KO^* mice were viable and exhibited no gross morphological defects. Consistent with prior findings, *Znrf3^KO/KO^* mice were embryonic lethal and displayed the expected lens malformation phenotype in all embryos (Fig. 1A).

**Figure 1.**
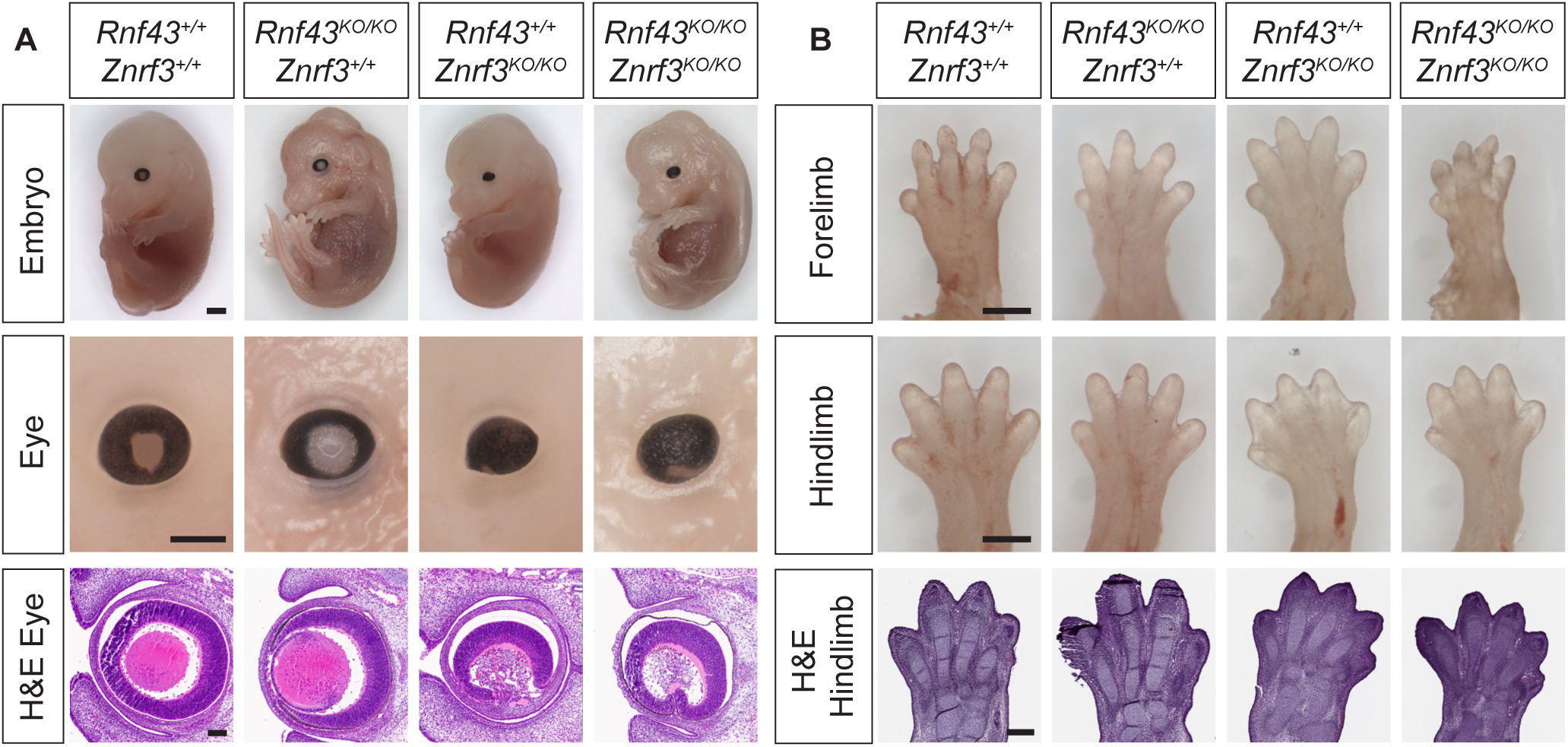
Germline deletion of *Znrf3*, alone or in combination with *Rnf43*, disrupts eye development without inducing limb duplication. (A) Representative whole-mount images of E16.5 control, germline *Rnf43^KO/KO^*, *Znrf3^KO/KO^*, and double knockout embryos. Higher-magnification views of the eye and corresponding hematoxylin and eosin (H&E)–stained eye sections are shown. Embryos lacking *Znrf3*, either alone or in combination with *Rnf43*, exhibit abnormal eye morphology. Embryo scale bar = 1 mm; eye whole-mount scale bar = 500 µm; H&E eye section scale bar = 100 µm. (B) Representative whole-mount images of E16.5 forelimbs and hindlimbs from control, *Rnf43^KO/KO^*, *Znrf3^KO/KO^*, and double knockout embryos. No supernumerary limbs or gross abnormalities in limb patterning were observed. H&E-stained hindlimb sections are shown to assess tissue morphology. Forelimb and hindlimb scale bars = 500 µm; H&E hindlimb section scale bar = 250 µm.

We next intercrossed single knockouts to generate compound knockout mice. *Rnf43^KO/KO^; Znrf3^KO/+^* mice were viable, fertile, and phenotypically normal. Timed matings from this genotype yielded *Rnf43^KO/KO^; Znrf3^KO/KO^* double knockout embryos, which were consistently recovered up to embryonic day (E)16.5. These embryos exhibited defective lens development resembling that observed in *Znrf3^KO/KO^* embryos (Fig. 1A).

Despite previous reports of limb duplication in *Xenopus tropicalis* following simultaneous loss of *rnf43* and *znrf3*^14^, no evidence of supernumerary limb formation was observed in *Rnf43^KO/KO^; Znrf3^KO/KO^* mouse embryos (Fig. 1A). Histology of E16.5 fore- and hindlimbs revealed no gross patterning defects (Fig. 1B). Thus, combined germline deletion of *Rnf43* and *Znrf3* in mice does not recapitulate the supernumerary limb phenotype observed in amphibians.

### Znrf3 Is the Predominantly Expressed Paralog in Mature Osteoblasts

To assess the relative expression of *Rnf43* and *Znrf3* in the osteoblast lineage, we analyzed transcript levels in calvarial osteoblasts derived from *Ocn*-Cre; *mTmG* mice. The *mTmG* allele is a Cre-dependent reporter that expresses membrane-targeted GFP (mG) following Cre-mediated excision of a loxP-flanked Tomato (mT) cassette, thereby marking cells with *Ocn*-driven recombination. Calvarial bones were collected from neonatal *Ocn*-Cre*; mTmG* mice, enzymatically digested into single-cell suspensions, and GFP⁺ osteoblasts were isolated by flow cytometry.

Bulk RNA-seq analysis of sorted osteoblasts revealed that both *Rnf43* and *Znrf3* are expressed in mature osteoblasts, though at markedly different levels. *Rnf43* transcripts were detected at log2(TPM) values ranging from 7.59 to 8.01 across biological replicates, while *Znrf3* expression ranged from 10.68 to 10.88 (Fig. 2), corresponding to ∼8-fold higher expression of Znrf3 than Rnf43 in mature osteoblasts. Notably, both genes exhibited low inter-sample variability, consistent with stable and reproducible expression in the osteoblast population targeted by *Ocn*-Cre.

**Figure 2.**
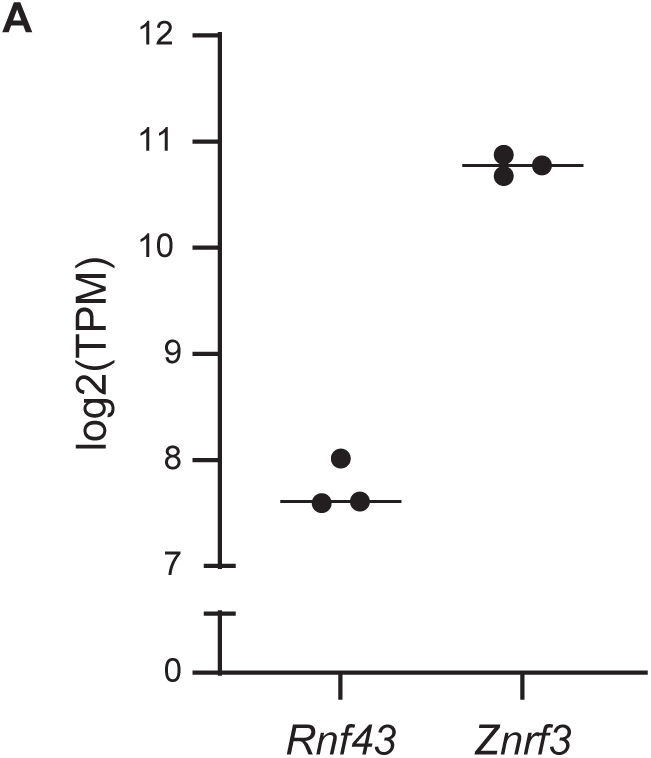
Expression levels of *Rnf43* and *Znrf3* in *Ocn*⁺ osteoblasts. (A) GFP⁺ osteoblasts were isolated by FACS from calvarial digests of *Ocn*-Cre*; mTmG* neonatal mice. RNA was extracted from sorted cells and analyzed by bulk RNA sequencing. Transcript abundance was quantified using TPM (Transcripts Per Million) and log₂-transformed. Boxplots show that both *Rnf43* and *Znrf3* are expressed in osteoblasts, with *Znrf3* exhibiting significantly higher expression (∼3 log₂ units, ∼8-fold difference) across biological replicates.

These data confirm that *Znrf3* is the predominant paralog expressed in mature osteoblasts and suggests it may play a more prominent role than *Rnf43* in regulating Wnt signaling within this cell type.

### Osteoblast-Specific Deletion of Znrf3, but Not Rnf43, Alters Skeletal Architecture Trabecular Bone Analysis

To determine the individual contributions of the *Rnf43/Znrf3* paralogs in osteoblast-mediated bone homeostasis, we generated mice carrying osteoblast-specific deletions of either *Znrf3* or *Rnf43* using *Osteocalcin (Ocn)-Cre*. Because both genes encode transmembrane E3 ubiquitin ligases that negatively regulate Wnt/β-catenin signaling, we hypothesized that deletion of either paralog would increase bone mass. This expectation was based on our prior studies demonstrating that *Ocn*-Cre-mediated deletion of *Adenomatous Polyposis Coli (Apc)*, a central negative regulator of β-catenin, produces a robust high-bone-mass phenotype ^16^.

Micro-computed tomography (µCT) analysis of femurs from *Znrf3^cKO/cKO^* mice at 3 and 6 months of age revealed an unexpected trabecular bone phenotype. Representative three-dimensional reconstructions of distal femoral trabecular bone are shown in Fig. 3A, with visible architectural differences in 6-month-old animals. Quantitative analysis revealed no consistent differences in trabecular bone mineral density (BMD) at 3 months in either sex; however, 6-month-old male knockout mice exhibited reduced BMD relative to controls (Fig. 3B). Similarly, bone volume fraction (BV/TV), which quantifies the fraction of mineralized bone within total tissue volume, was significantly decreased in 6-month-old male knockout mice (Fig. 3C). Trabecular thickness (Tb.Th) was not significantly altered (Fig. 3D), whereas trabecular separation (Tb.Sp) was increased and trabecular number (Tb.N) was decreased in 6-month-old males (Fig. 3E, F), indicating reduced trabecular connectivity and overall bone mass. Together, these findings demonstrate age- and sex-dependent reductions in trabecular bone mass following osteoblast-specific deletion of *Znrf3*, with the most pronounced effects observed in 6-month-old male mice.

**Figure 3.**
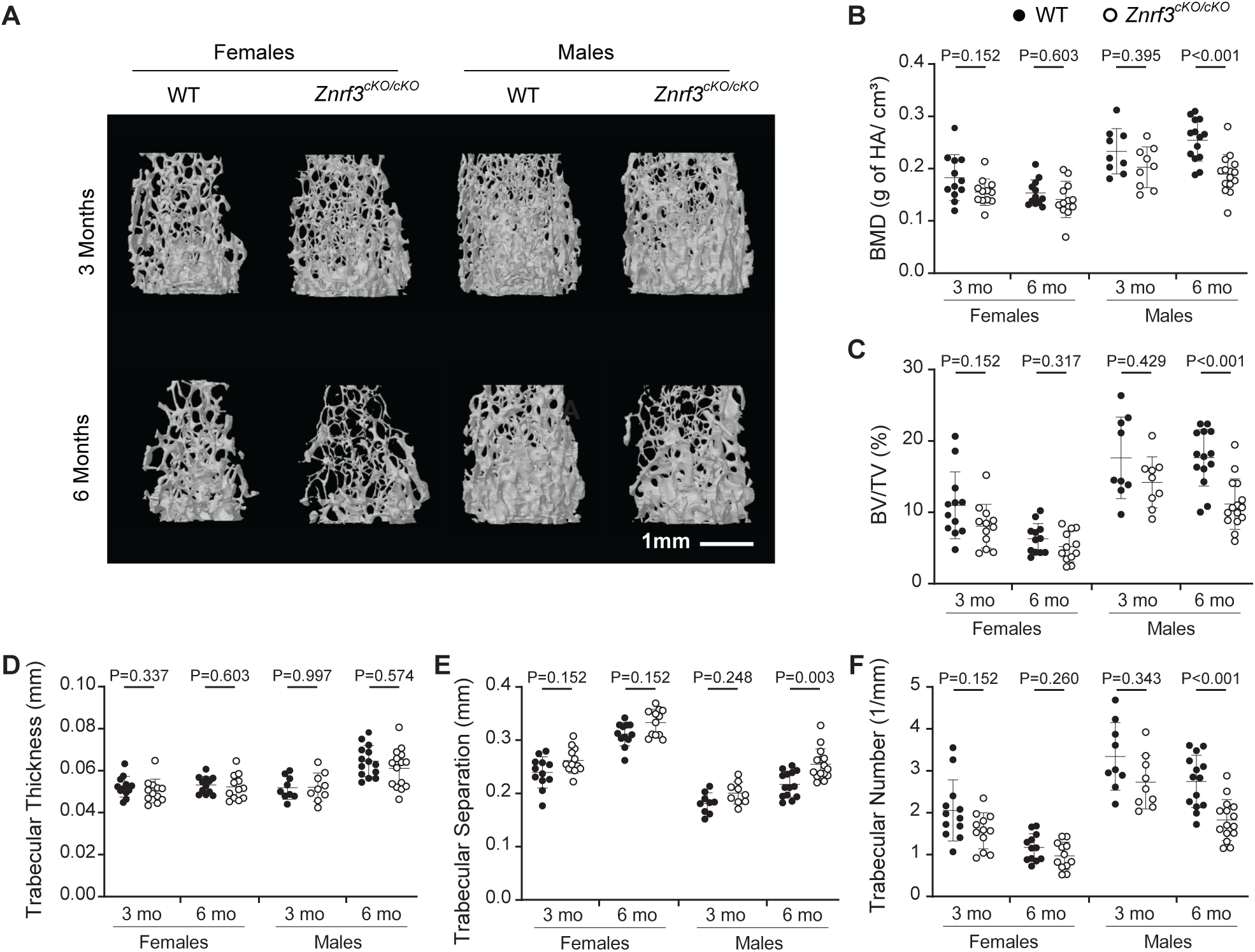
Osteoblast-specific deletion of *Znrf3* alters trabecular bone microarchitecture. (A) Representative micro-computed tomography (µCT) images of distal femoral trabecular bone from 3- and 6-month-old female and male control and *Znrf3^cKO/cKO^* mice. Trabecular parameters quantified by µCT include (B) bone mineral density (BMD; g hydroxyapatite/cm³), (C) percent bone volume fraction (BV/TV), (D) trabecular thickness (Tb.Th), (E) trabecular separation (Tb.Sp), and (F) trabecular number (Tb.N). Sample sizes were as follows: 3-month females (control and knockout n = 12 per group), males (control and knockout n = 9 per group) and 6-month females (control and knockout n = 12 per group) and males (control n = 14; knockout n = 15). Data are presented as dot plots with individual data points shown. The horizontal line indicates the group mean, and error bars represent standard deviation (SD). Statistical analysis was performed using robust linear regression on log-transformed outcome variables. P values are indicated on each plot.

To further investigate potential cellular mechanisms underlying the trabecular phenotype, distal femoral sections from 3- and 6-month-old male *Znrf3^cKO/cKO^* mice were analyzed by static histomorphometry. Trabecular sections were stained with Goldner’s Trichrome to visualize and quantify adipocytes, osteoblasts, and osteoid matrix. Parameters associated with bone formation and marrow composition, including bone volume/tissue volume (BV/TV), adipocyte volume/tissue volume (Ad.V/TV), osteoid volume/bone volume (OV/BV), osteoid surface/bone surface (OS/BS), osteoid width (O.Wi), and osteoblast number/bone surface (N.Ob/BS), were not significantly different from controls (Supplemental Table 1). These findings indicate that although structural deficits in trabecular bone were detected by µCT, static histomorphometry did not reveal overt changes in osteoblast number, osteoid production, or marrow adiposity.

### Cortical Bone Analysis

Analysis of femoral midshaft cortical bone further revealed changes in bone geometry. Representative cross-sectional images (Fig. 4A) demonstrate increased cortical cross-sectional size accompanied by reduced cortical thickness in knockout mice. Quantitative analysis showed no consistent differences in tissue mineral density (TMD) between genotypes (Fig. 4B), indicating preserved mineralization. In contrast, total tissue area (T.Ar) was increased in both sexes, particularly at 6 months of age (Fig. 4C), suggesting expansion of the cortical envelope. Bone area (B.Ar) followed a similar trend (Fig. 4D) but did not reach statistical significance.

**Figure 4.**
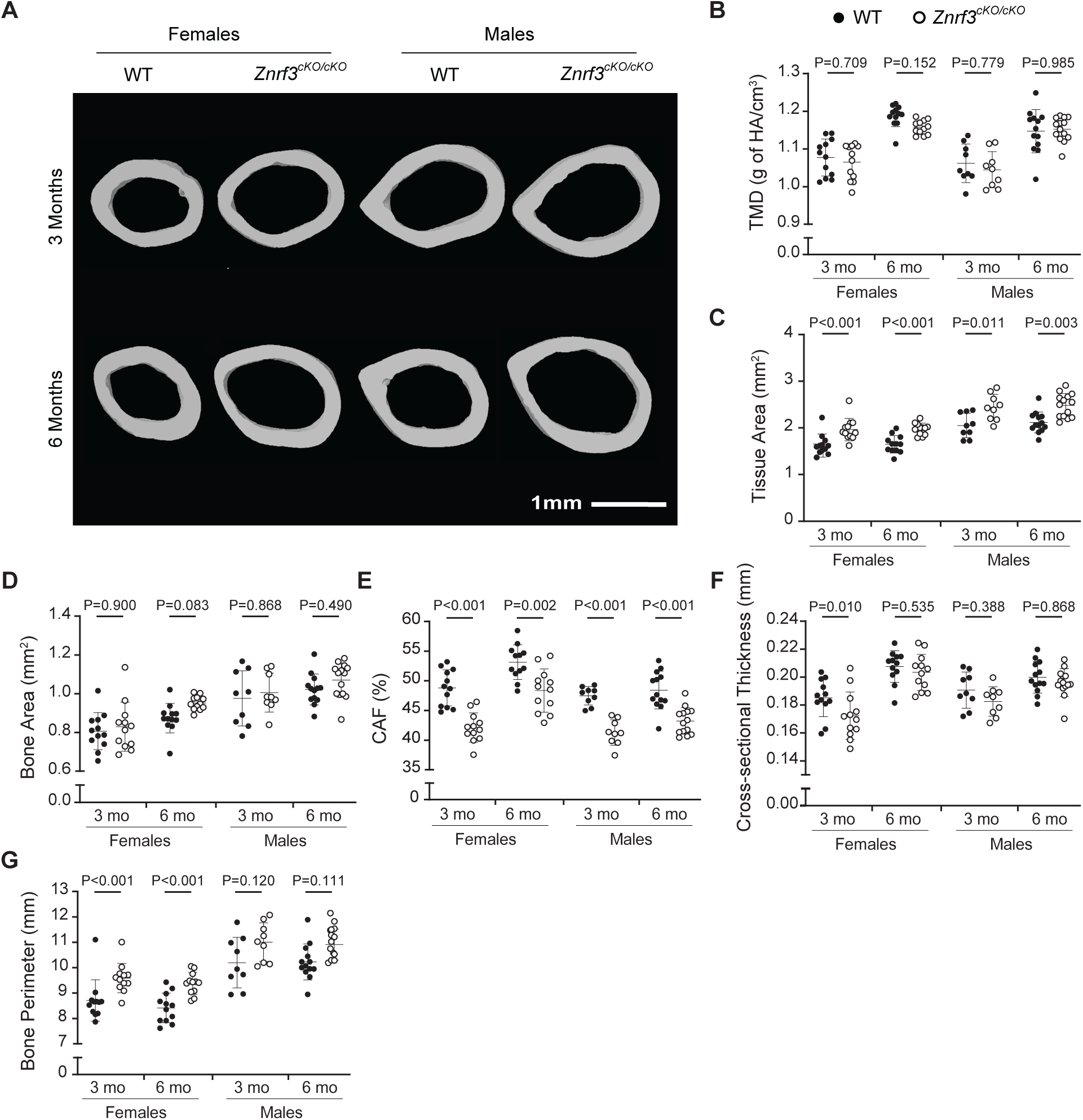
Osteoblast-specific deletion of *Znrf3* alters cortical bone geometry at the femoral midshaft. (A) Representative micro-computed tomography (µCT) images of femoral midshaft cortical bone from 3- and 6-month-old female and male control and *Znrf3^cKO/cKO^* mice. Cortical parameters quantified by µCT include (B) tissue mineral density (TMD; g hydroxyapatite/cm³), (C) total tissue area (T.Ar), (D) bone area (B.Ar), (E) cortical area fraction (CAF), (F) cross-sectional thickness (Cs.Th), and (G) bone perimeter (B.Pm). Sample sizes were as follows: 3-month females (control and knockout n = 12 per group), males (control and knockout n = 9 per group) and 6-month females (control and knockout n = 12 per group) and males (control n = 14; knockout n = 15). Data are presented as dot plots with individual data points shown. The horizontal line indicates the group mean, and error bars represent standard deviation (SD). Statistical analysis was performed using robust linear regression on log-transformed outcome variables. P values are indicated on each plot.

Despite this increase in cross-sectional size, cortical area fraction (CAF) was reduced in knockout mice (Fig. 4E). Cross-sectional thickness (Cs.Th) was significantly reduced in 3-month female knockout animals (Fig. 4F). Bone perimeter (B.Pm) was increased in female knockout animals (Fig. 4G) and trended upward in males, supporting outward expansion of the diaphyseal envelope without increased cortical mass.

To determine whether altered cortical geometry was associated with changes in bone formation dynamics, fluorochrome labeling was performed in 6-month-old male *Znrf3^cKO/cKO^* mice. Animals received fluorescent labels at 4, 10, 24, and 25 weeks of age. Mineral apposition rate (MAR) and bone formation rate (BFR) were quantified in femoral midshaft cortical sections across the entire cortex and separately within anterior and posterior regions.

MAR and BFR were calculated for both endocortical and periosteal surfaces over three labeling intervals (4–10 weeks, 10–25 weeks, and 4–25 weeks). When analyzed across the whole cortical surface, no significant differences were detected in endocortical MAR, endocortical BFR, periosteal MAR, or periosteal BFR between *Znrf3^cKO/cKO^* and control mice (Supplemental Table 2). To assess whether localized effects were obscured by whole-bone averaging, MAR and BFR were additionally analyzed in anterior and posterior cortical regions independently.

Again, no significant differences were observed in endocortical or periosteal MAR or BFR on either surface in knockout mice compared to controls (Supplemental Table 2). These findings indicate that the cortical geometric changes observed following *Znrf3* deletion are not accompanied by detectable alterations in steady-state mineral apposition or bone formation rates at 6 months of age.

In contrast to the phenotype observed following deletion of *Znrf3*, osteoblast-specific deletion of *Rnf43* did not result in detectable changes in trabecular or cortical bone parameters. µCT analysis of *Rnf43^cKO/cKO^* mice at 3 and 6 months of age revealed no significant differences in trabecular BMD, BV/TV, trabecular thickness, number, or separation relative to controls.

Likewise, cortical bone parameters including tissue area, bone area, cortical thickness, cortical area fraction, bone perimeter, and tissue mineral density were comparable between genotypes across sexes and time points (Supplemental Tables 3-4).

Together, these data demonstrate that osteoblast-specific loss of *Znrf3*, but not *Rnf43*, disrupts trabecular bone mass and cortical geometry, despite their shared roles as negative regulators of Wnt/β-catenin signaling. These findings identify *Znrf3* as the dominant functional paralog regulating skeletal architecture in mature osteoblasts.

### Osteoblast-Specific Deletion of *Rnf43* and *Znrf3* Does Not Exacerbate the *Znrf3* Conditional Knockout Skeletal Phenotype

Previous studies in other tissues, including the intestinal epithelium, have demonstrated functional redundancy between *Rnf43* and *Znrf3*, such that simultaneous loss of both genes produces more severe phenotypes than deletion of either gene alone ^11, 17–19^. Although osteoblast-specific deletion of *Rnf43* did not produce a detectable skeletal phenotype, we next examined whether combined deletion of *Rnf43* and *Znrf3* in mature osteoblasts would exacerbate the abnormalities observed following loss of *Znrf3* alone.

Micro-CT analysis of femurs from double conditional knockout mice at 3 and 6 months of age revealed trabecular bone phenotypes comparable to those observed in *Znrf3* single knockout mice. Representative three-dimensional reconstructions of distal femoral trabecular bone are shown in Fig. 5A, where 6-month-old male double knockout mice display visibly reduced trabecular density relative to controls. Quantitative analysis demonstrated reduced trabecular bone mineral density (BMD) in 6-month-old male double knockout mice (Fig. 5B), accompanied by reduced bone volume fraction (BV/TV) (Fig. 5C). Trabecular thickness (Tb.Th) was not significantly altered, although a modest trend toward reduction was observed in 6-month-old females and males (Fig. 5D). Consistent with the *Znrf3* single knockout phenotype, 6-month-old male double knockout mice exhibited increased trabecular separation (Tb.Sp) and decreased trabecular number (Tb.N) (Fig. 5E, F), further supporting disruption of trabecular microarchitecture.

**Figure 5.**
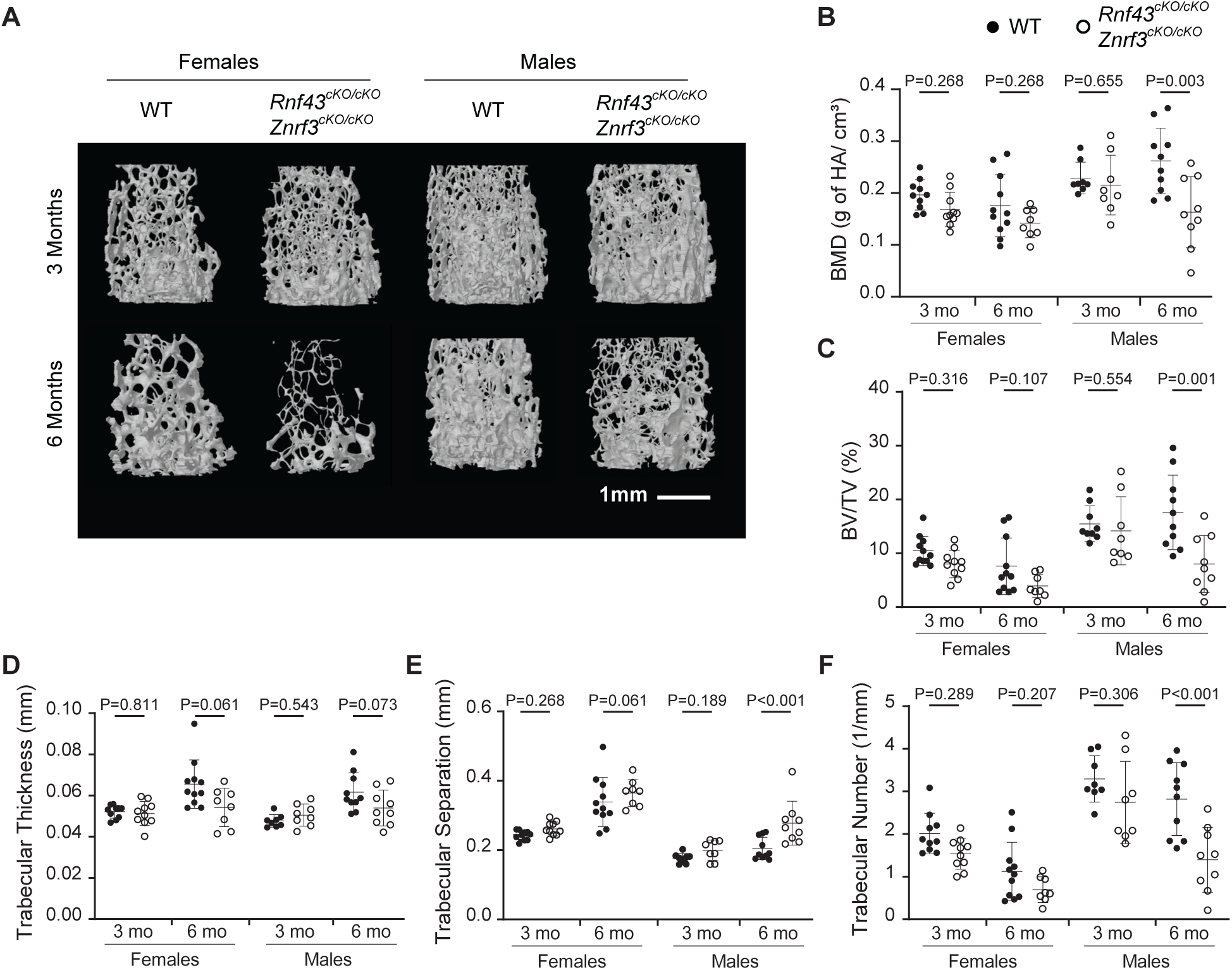
Osteoblast-specific deletion of *Rnf43* and *Znrf3* does not exacerbate trabecular bone defects. (A) Representative micro-computed tomography (µCT) images of distal femoral trabecular bone from 3- and 6-month-old female and male control and *Rnf43^cKO/cKO^; Znrf3^cKO/cKO^* mice. Trabecular parameters quantified by µCT include (B) bone mineral density (BMD; g hydroxyapatite/cm³), (C) percent bone volume fraction (BV/TV), (D) trabecular thickness (Tb.Th), (E) trabecular separation (Tb.Sp), and (F) trabecular number (Tb.N). Sample sizes were as follows: 3-month females (control and knockout n = 10 per group), males (control and knockout n = 8 per group) and 6-month females (control n = 11; knockout n = 8) and males (control n = 10; knockout n = 9). Data are presented as dot plots with individual data points shown. The horizontal line indicates the group mean, and error bars represent standard deviation (SD). Statistical analysis was performed using robust linear regression on log-transformed outcome variables. P values are indicated on each plot.

Static histomorphometric analysis of distal femoral trabecular bone from 6-month-old male double knockout mice did not reveal significant differences in BV/TV, osteoid parameters, osteoblast number, or marrow adiposity compared to controls (Supplemental Table 1). Thus, although µCT detected structural deficits, static indices did not identify overt cellular alterations in these animals. Importantly, comparison of double *Rnf43*/*Znrf3* knockout mice with *Znrf3* single knockout mice did not reveal additional worsening of any trabecular parameter, indicating that deletion of *Rnf43* does not exacerbate the trabecular defects caused by loss of *Znrf3*.

To determine whether the cortical phenotype was similarly affected, femoral midshaft cortical bone from control and double knockout mice was analyzed by µCT (Fig. 6A). Tissue mineral density (TMD) did not differ between genotypes at either age (Fig. 6B), consistent with findings in *Znrf3* single knockout mice. Total tissue area (T.Ar) was increased in male double knockout mice at both 3 and 6 months (Fig. 6C), whereas females showed a significant increase only at 3 months. Bone area (B.Ar) was increased exclusively in 3-month-old female double knockout mice compared to controls (Fig. 6D). Although cortical area fraction (CAF) was reduced (Fig. 6E), cross-sectional thickness (Cs.Th) was not significantly altered (Fig. 6F). Bone perimeter (B.Pm) was increased in male double knockout mice at both ages and in females at 3 months (Fig. 6G), supporting outward expansion of the diaphyseal envelope without increased cortical thickness. As with trabecular bone, comparison of double knockout and *Znrf3* single knockout mice revealed no additional worsening of cortical parameters.

**Figure 6.**
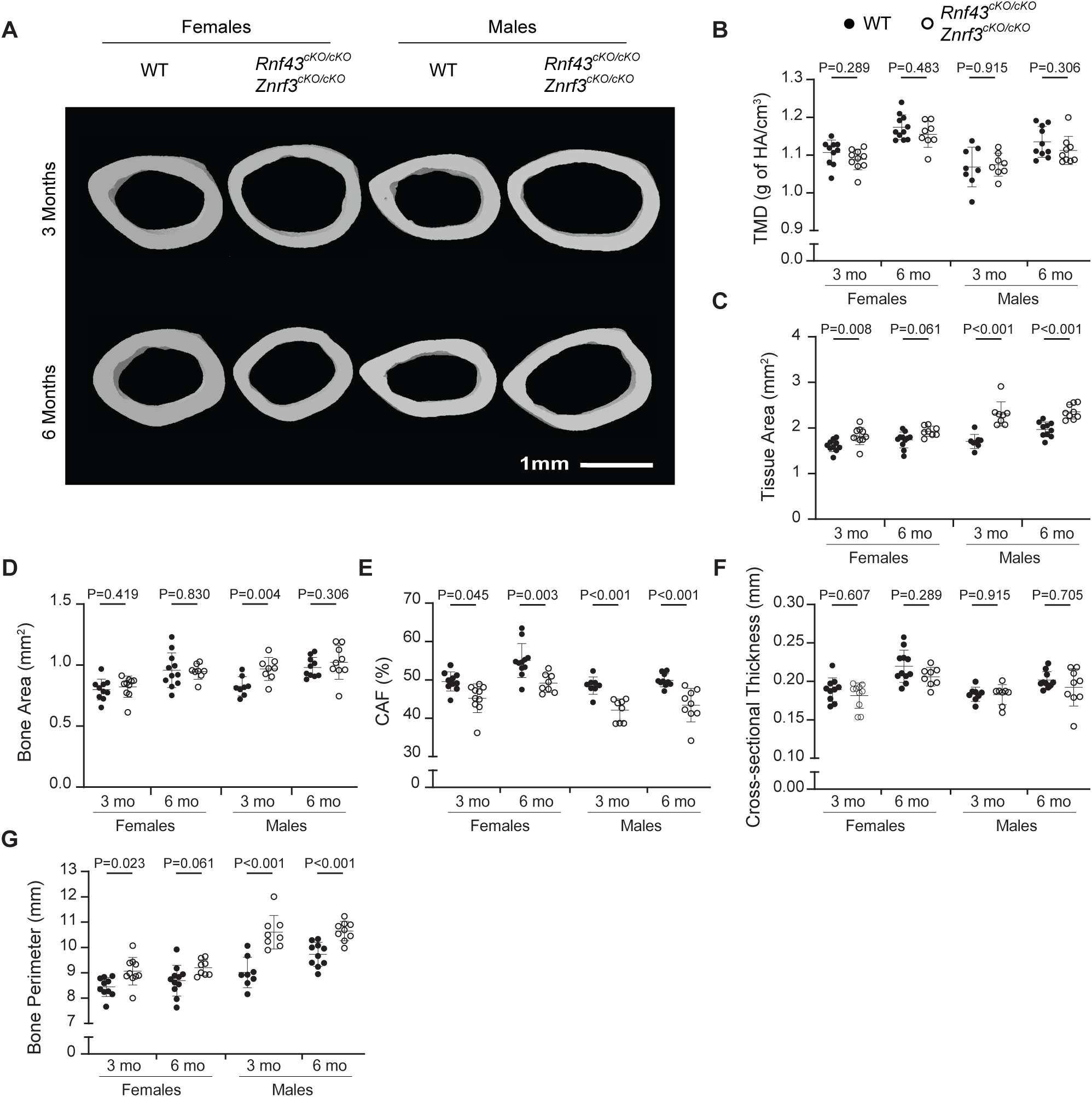
Cortical bone parameters in osteoblast-specific *Rnf43* and *Znrf3* knockout mice. (A) Representative micro-computed tomography (µCT) images of femoral midshaft cortical bone from 3- and 6-month-old female and male control and *Rnf43^cKO/cKO^; Znrf3^cKO/cKO^*mice. Cortical parameters quantified by µCT include (B) tissue mineral density (TMD; g hydroxyapatite/cm³), (C) total tissue area (T.Ar), (D) bone area (B.Ar), (E) cortical area fraction (CAF), (F) cross-sectional thickness (Cs.Th), and (G) bone perimeter (B.Pm). Sample sizes were as follows: 3-month females (control and knockout n = 10 per group), males (control and knockout n = 8 per group) and 6-month females (control n = 11; knockout n = 8) and males (control n = 10; knockout n = 9). Data are presented as dot plots with individual data points shown. The horizontal line indicates the group mean, and error bars represent standard deviation (SD). Statistical analysis was performed using robust linear regression on log-transformed outcome variables. P values are indicated on each plot.

Together, these data demonstrate that simultaneous loss of *Rnf43* and *Znrf3* in mature osteoblasts does not enhance the skeletal phenotype observed following loss of *Znrf3* alone. These findings further support *Znrf3* as the dominant functional paralog regulating trabecular bone mass and cortical geometry in the osteoblast lineage, with limited compensatory contribution from *Rnf43* in this context.

## DISCUSSION

In this study, we define a lineage-specific and largely non-redundant role for the Wnt receptor E3 ligase ZNRF3 in regulating skeletal architecture. Although RNF43 and ZNRF3 are frequently regarded as functionally redundant negative regulators of Wnt signaling, our data demonstrate that this assumption does not fully extend to the mature osteoblast lineage. Instead, *Znrf3* emerges as the dominant paralog in bone, where its deletion produces architectural remodeling rather than the high-bone-mass phenotype predicted from canonical Wnt gain-of-function models. These findings refine our understanding of how receptor-level control of Wnt signaling shapes skeletal biology and highlight the context-dependent nature of Wnt pathway regulation *in vivo*.

### Species and developmental context shape Rnf43/Znrf3 function

Simultaneous disruption of *rnf43* and *znrf3* in *Xenopus* produces dramatic limb duplication, implicating these ligases in early limb patterning. In those studies, gene disruption occurred during early embryogenesis, affecting limb field specification. In contrast, germline deletion of both genes in mice did not induce supernumerary limb formation. This divergence likely reflects differences in developmental timing and experimental context rather than simple species divergence. Whereas Xenopus models interrogate early patterning programs, our mammalian analyses examined either germline deletion or lineage-restricted deletion in mature osteoblasts, contexts in which limb specification is already established. These findings underscore the importance of developmental stage and tissue context when interpreting the skeletal consequences of Wnt pathway perturbation.

### Receptor-level modulation is not equivalent to β-catenin stabilization

A central and unexpected finding of this study is that osteoblast-specific deletion of *Znrf3* does not phenocopy canonical Wnt gain-of-function models. Stabilization of β-catenin or deletion of *Apc* in osteoblasts produces dramatic increases in bone mass ^4, 20, 21^. In contrast, *Znrf3* deletion resulted in age- and sex-dependent reductions in trabecular bone mass, most prominently in 6-month-old male mice, along with altered cortical geometry.

This distinction highlights an important conceptual point: receptor-level regulation of Wnt signaling is not functionally equivalent to downstream stabilization of β-catenin. *Apc* deletion bypasses extracellular ligand availability and feedback inhibition, driving sustained and cell-autonomous activation of canonical transcriptional programs. In contrast, ZNRF3 regulates Frizzled receptor abundance at the plasma membrane, thereby modulating responsiveness to local Wnt ligands. The skeletal consequences of *Znrf3* deletion therefore remain constrained by ligand availability, receptor composition, feedback inhibitors, and stage-specific signaling thresholds. These findings suggest that modulation of receptor turnover may recalibrate Wnt sensitivity rather than uniformly amplify canonical output.

### Architectural remodeling rather than net bone gain

The trabecular phenotype following *Znrf3* deletion is characterized by reduced bone mineral density, decreased bone volume fraction, reduced trabecular number, and increased trabecular separation, without consistent changes in trabecular thickness. This pattern indicates loss of trabecular connectivity rather than uniform thinning.

In the cortical compartment, *Znrf3* deletion resulted in increased cross-sectional size accompanied by reduced cortical area fraction and altered geometry, while tissue mineral density remained unchanged. Cross-sectional thickness was only selectively reduced in certain sex- and age-specific contexts. Together, these findings indicate redistribution of mineralized tissue rather than uniform cortical thinning or global changes in mineralization.

Although we did not assess mechanical properties, such geometric remodeling suggests altered spatial organization of bone rather than simple changes in total bone mass. The age- and sex-dependent nature of these effects further indicates that receptor-level Wnt regulation interacts with systemic or hormonal factors to shape skeletal architecture.

### Potential balance between canonical and noncanonical signaling

ZNRF3 targets multiple Frizzled receptors and therefore has the capacity to influence both canonical and noncanonical Wnt pathways. While bone biology has historically emphasized β-catenin–dependent signaling as a primary anabolic driver, noncanonical pathways also contribute to osteoblast maturation, cytoskeletal organization, and osteoblast–osteoclast coupling.

One possibility is that loss of *Znrf3* alters the balance between canonical and noncanonical Wnt signaling rather than uniformly increasing β-catenin activity. Because pathway activity was not directly measured, the relative contributions of these branches remain undefined. However, altered signaling balance may contribute to the architectural redistribution observed here. More broadly, these findings reinforce that the skeletal consequences of perturbing Wnt regulators cannot be predicted solely by their canonical pathway annotation.

### Limited redundancy in mature osteoblasts

Although RNF43 and ZNRF3 function redundantly in highly proliferative tissues such as the intestinal epithelium, simultaneous deletion of both genes in mature osteoblasts did not exacerbate the *Znrf3* single-mutant phenotype. Bulk RNA-seq analysis demonstrated substantially higher expression of *Znrf3* relative to *Rnf43* in *Ocn*⁺ osteoblasts, suggesting that paralog dominance may be dictated by lineage-specific transcriptional programs. These data reinforce that genetic redundancy is context-dependent and cannot be inferred solely from shared biochemical activity.

### Implications for therapeutic modulation of the RSPO–Wnt axis

R-spondin proteins potentiate Wnt signaling in part by inhibiting RNF43 and ZNRF3, and biologics targeting this axis are under development for regenerative medicine and oncology. Our findings suggest that sustained modulation of receptor-level Wnt regulators in bone may influence skeletal architecture in ways not fully predicted by canonical β-catenin activation models. Alterations in cortical geometry without proportional increases in bone mass emphasize the importance of evaluating structural outcomes, not solely bone density, when therapeutically targeting upstream Wnt modulators.

### Limitations and future directions

This study focused on mature osteoblasts using *Ocn*-Cre and therefore does not address potential roles for *Rnf43* and *Znrf3* in earlier mesenchymal progenitors or osteocytes. Future studies examining stage-specific deletions will be necessary to fully define how receptor-level Wnt regulation shapes skeletal function.

## Conclusion

Together, our findings reveal that ZNRF3 plays a dominant and largely non-redundant role in regulating skeletal architecture within the mature osteoblast lineage. Contrary to expectations derived from canonical Wnt activation models, receptor-level disruption of Wnt regulation does not produce uniform anabolic effects but instead remodels bone geometry and reduces trabecular mass in an age- and sex-dependent manner. These results underscore the importance of cellular context, ligand environment, and pathway balance in determining the skeletal consequences of Wnt perturbation and refine our understanding of how upstream Wnt regulators shape tissue architecture *in vivo*.

## MATERIALS AND METHODS

### Mouse Strains and Genotyping

All mice were maintained following institutional animal care and use guidelines, and experimental protocols were approved by the Institutional Animal Care and Use Committee of the Van Andel Institute.

*Ocn*-Cre expressing mice are used extensively by our laboratory ^22–25^ and are on an FVBN background. Mouse strains containing floxed alleles of either *Rnf43* or *Znrf3* were obtained from Hans Clevers ^11^ and were on a mixed genetic background of 129/SvJ and C57BL/6J.

To create mice in which Rnf43 and/or Znrf3 were globally deleted (germline deletion), we crossed the *Rnf43^Flox^* and *Znrf3^Flox^* strains to a Cre strain which is under the control of a human cytomegalovirus minimal promoter (CMV-Cre) (Jackson Labs, 006054) ^26^. The resulting mice were backcrossed to C57BL/6J to remove the CMV-Cre transgene before heterozygous intercrosses. Rnf43^KO/KO^; Znrf3^KO/+^ mice were viable and fertile and used for timed matings to generate embryos lacking both genes globally (Rnf43^KO/KO^; Znrf3^KO/KO^).

We used the following primers to genotype the Rnf43^flox^ and Znrf3^flox^ lines: Rnf43-loxP-Fwd (GAAGCAGACAATGAAGCGAAT) and Rnf43-loxP-Rev (TAGTGCCCCACAGAGGACA) which amplify a 345 bp flox and a 284 bp wild-type product. Primers Znrf3-loxP-Fwd (CACACCCTGACCCTACGAA) and Znrf3-loxP-Rev (TTACCACACCCATACCCAACT) amplify a 405 bp flox and a 324 bp wild-type product. After Cre-mediated recombination, we used the following primers to determine Rnf43 and Znrf3 knockout animals: Rnf43-I6-Fwd (CACCGCCATTACTCTCTATTCC), Rnf43-loxP-Fwd (see above), and Rnf43-I7-Rev (GAGCAGTCGGTGTTCTTAACT) amplify a 744 bp flox, a 621 bp wild-type, and a 274 bp knockout product. Znrf3-loxP-Fwd (see above), Znrf3-AS-Fwd (AGTTAGGGAAGAGAGTGGGAA), and Znrf3-loxP-Rev (see above) amplify a 403 bp flox, a 324 bp wild-type, and a 247 bp knockout product.

### Histology of germline deleted embryos

Timed matings of intercrossed Rnf43^KO/KO^; Znrf3^KO/+^ were established in the evening, and the following morning females were assessed for vaginal plugs (E0.5). Pregnant females were euthanized, and embryos were harvested at E16.5. Embryos were fixed in 10% NBF for 24 hours at room temperature, followed by dehydration and paraffin embedding. Sagittal sections (5µm) were cut and stained for hematoxylin and eosin. Slides were scanned (Zeiss AxioScan.Z1), and cross-sections were assessed for phenotypic changes.

### Microcomputed Tomography (µCT)

For skeletal analysis, femurs were isolated from 3 and 6-month-old female and male mice, fixed in 10% neutral-buffered formalin (NBF) at room temperature for 48 h, and then changed to 70% ethanol before analysis.

These samples were analyzed using a SkyScan 1172 µCT system (Bruker MicroCT: Kontich, Belgium) ^28^. Femora were scanned in 70% ethanol using an X-ray voltage of 60 kV, current of 167 µA, and 0.5 mm aluminum filter with a voxel size of 7.98 µm. Femoral images were reconstructed using NRecon 1.7.4.6 (Bruker MicroCT). The mineralized tissue was oriented, and a volume of interest (VOI) was defined using DataViewer 1.5.6.3 (Bruker MicroCT).

Regions of interest (ROI) were defined for cortical and trabecular bone using CTAn 1.18.8.0 (Bruker MicroCT). A trabecular ROI was drawn in the distal epiphysis for each femur, starting 0.25 mm proximal to the growth plate and 2.5 mm in height. To define each cortical ROI’s position, 45% of the femur length was calculated and used to set the distal end of the region. The ROI was 0.8 mm in height toward the proximal end of the bone, within the midshaft. Trabecular 3D analysis was performed to quantify bone mineral density (BMD), bone volume/tissue volume (BV/TV), bone surface/bone volume, trabecular thickness (Tb.Th), trabecular separation (Tb.Sp), and trabecular number (Tb.N). Cortical 2D analysis was performed to quantify tissue mineral density (TMD), tissue area (T.Ar), bone area (B.Ar), cortical area fraction (bone area/tissue area, CAF), cross-sectional thickness (Cs.Th), and bone perimeter (B.Pm).

### Fluorochrome labeling

Male mice were injected with fluorochrome over 6 months to measure femoral bone formation indices. Mice were injected (mg of fluorochrome/kg body weight) with 10 mg/kg of calcein (Sigma, C0875) at 4 weeks of age, 20 mg/kg of alizarin complexone (Sigma, A3882) at 10 weeks of age, 75 mg/kg of demeclocycline (Sigma, D6140) at 24 weeks of age, and again with 10 mg/kg of calcein at 25 weeks of age. All dosages were split between intraperitoneal and subcutaneous injections. At 26 weeks of age, animals were sacrificed, femurs were isolated, fixed in NBF for 48 hours, and stored in 70% ethanol in the dark.

### Histomorphometry

Static and dynamic histomorphometry was performed on femurs from 3- and 6-month-old mice. Detailed steps for preparation and processing are available in ^28^. In brief, femoral samples were fixed in NBF at room temperature for 48 hours and kept in 70% ethanol before processing.

Femurs were dehydrated in graded ethanol in 2-4 hour increments and cleared using xylene. Samples were infiltrated with 85-100% MMA and an 15% dibutyl phthalate over several days. Femurs were coronally sectioned for static analysis and cross-sectioned for dynamic analysis. All analysis was performed using Bioquant Osteo v19.2.60.

Static histomorphometry was performed on coronally sectioned femurs. The samples were deplasticized and stained following the Goldner’s Trichrome protocol. Slides were imaged using the Leica Aperio AT2 at 20X magnification. The regions of interest (ROI) drawn were 2000 µm long starting from the distal femoral growth plate. The edges of the ROI were kept 200 µm from the growth plate and all cortical bone. The parameters measured included bone volume/tissue volume (BV/TV), adipocyte volume/tissue volume (Ad.V/TV), osteoid volume/bone volume (OV/BV), osteoid surface/bone surface (OS/BS), osteoid width (O.Wi), and number of osteoblasts/bone surface (N.Ob/BS).

Dynamic histomorphometry was performed on femoral cross-sections taken from the midshaft of 6-month-old, fluorochrome injected, male mice. The samples were cover slipped and imaged using a Zeiss Axio Imager.A2 fluorescence microscope and Lumen Dynamics X-Cite Series 120 Q, QImaging QIClick Digital CCD camera, under an ET – DAPI/FITC/Texas Red (Chroma product # 69002) filter, at 20X magnification. The analysis was separated into three regions of the femur: whole bone, anterior, and posterior. The parameters measured for each included endocortical mineral apposition rate (MAR), endocortical bone formation rate (BFR), periosteal MAR, and periosteal BFR.

### Statistics

All µCT and histomorphometry data were analyzed with robust linear regressions and log transformed outcomes. Robust regression was used because some influential outliers were identified based on Cook’s distance exceeding 4/n. We used robust regression to limit the influence of potential outliers on the estimation of the effects. These regressions used weights such that points with a large amount of influence are downweighted based on established methods. Here we specifically used MM-estimation, which works well under normality.

Benjamini-Hochberg false discovery rate corrections were used to adjust for multiple testing.

## Acknowledgements

We thank the following VAI core facilities for their assistance: Bioinformatics and Biostatistics (RRID:SCR_024762); Flow Cytometry (RRID:SCR_022685); Genomics (RRID:SCR_022913); Pathobiology and Biorepository (RRID:SCR_022912); Transgenics (RRID:SCR_022914); and Vivarium (RRID:SCR_023211). We also thank past and present members of the Williams Laboratory for their assistance and the Van Andel Institute for support.

