## Supplemental Files for "Deletion of the Wnt regulator Znrf3 alters bone geometry without inducing high bone mass"

**Supplemental Table 1. Static histomorphometric analysis of distal femoral trabecular bone in osteoblast-specific *Znrf3* knockout and double knockout mice.**

Coronal distal femoral sections were analyzed by static histomorphometry at 3 and 6 months of age. At 3 months, control (n = 5) and *Znrf3* knockout (n = 4) mice were examined. At 6 months, control (n = 12), *Znrf3* knockout (n = 10), double knockout control (n = 10), and double knockout (n = 8) mice were evaluated. Parameters measured included bone volume/tissue volume (BV/TV), adipocyte volume/tissue volume (Ad.V/TV), osteoid volume/bone volume (OV/BV), osteoid surface/bone surface (OS/BS), osteoid width (O.Wi), and osteoblast number per bone surface (N.Ob/BS; #/mm). Values are presented as mean ± SD.

| Parameter | 3 mo *Znrf3* Control | 3 mo *Znrf3* KO | 6 mo *Znrf3* Control | 6 mo *Znrf3* KO | 6 mo *Rnf43*/*Znrf3* Control | 6 mo *Rnf43*/*Znrf3* KO |
|---|---|---|---|---|---|---|
| **BV/TV (%)** | 15.0 ± 5.63 | 13.3 ± 4.04 | 15.6 ± 5.68 | 12.1 ± 3.59 | 11.53 ± 4.38 | 7.87 ± 3.62 |
| **Ad.V/TV (%)** | 1.05 ± 0.74 | 1.74 ± 0.93 | 1.26 ± 0.66 | 1.38 ± 0.66 | 1.68 ± 0.64 | 2.94 ± 1.23 |
| **OV/BV (%)** | 1.73 ± 0.91 | 2.21 ± 1.06 | 1.46 ± 0.67 | 1.29 ± 0.71 | 1.83 ± 0.57 | 2.53 ± 1.45 |
| **OS/BS (%)** | 12.5 ± 4.97 | 14.3 ± 6.81 | 12.1 ± 5.10 | 10.8 ± 3.94 | 14.3 ± 5.12 | 17.3 ± 10.7 |
| **O.Wi (µm)** | 2.53 ± 0.14 | 2.97 ± 0.54 | 2.73 ± 0.42 | 2.55 ± 0.44 | 2.58 ± 0.37 | 2.52 ± 0.34 |
| **N.Ob/BS (#/mm)** | 7.04 ± 2.38 | 9.38 ± 2.87 | 6.03 ± 2.17 | 6.39 ± 2.90 | 6.14 ± 2.69 | 6.75 ± 2.00 |

**Supplemental Table 2. Dynamic histomorphometric analysis of femoral midshaft cortical bone in fluorochrome-labeled osteoblast-specific *Znrf3* male mice.**

Male mice received fluorochrome injections at 4, 10, 24, and 25 weeks of age and were sacrificed one week after the final injection. Bone formation indices were quantified from femoral midshaft cortical cross-sections of control (n = 6) and *Znrf3* knockout (n = 8) mice. Mineral apposition rate (MAR) and bone formation rate (BFR) were calculated over three labeling intervals: 4-10 weeks (42 days), 10-25 weeks (105 days), and 4-25 weeks (147 days). Analyses were performed across the entire cortical surface and separately within anterior and posterior regions to assess regional differences in femoral growth.

| | | Endocortical MAR | | Endocortical BFR | | Periosteal MAR | | Periosteal BFR | |
|---|---|---|---|---|---|---|---|---|---|
| Time interval (days) | Region | Control | *Znrf3* KO | Control | *Znrf3* KO | Control | *Znrf3* KO | Control | *Znrf3* KO |
| 42 | Whole Bone | 1.878 ± 0.948 | 2.257 ± 1.192 | 0.872 ± 0.427 | 0.813 ± 0.495 | 2.096 ± 1.023 | 2.003 ± 0.370 | 1.086 ± 0.385 | 1.086 ± 0.385 |
| 42 | Anterior | 2.053 ± 0.985 | 2.245 ± 1.159 | 0.714 ± 0.334 | 0.783 ± 0.454 | 1.553 ± 0.844 | 1.287 ± 0.571 | 0.137 ± 0.090 | 0.238 ± 0.185 |
| 42 | Posterior | 0.357 ± 0.378 | 0.000 ± 0.000 | 0.044 ± 0.045 | 0.000 ± 0.000 | 2.152 ± 0.985 | 2.299 ± 0.599 | 0.524 ± 0.252 | 0.615 ± 0.169 |
| 105 | Whole Bone | 0.357 ± 0.194 | 0.312 ± 0.355 | 0.185 ± 0.092 | 0.112 ± 0.129 | 0.655 ± 0.321 | 0.927 ± 0.306 | 0.358 ± 0.195 | 0.586 ± 0.170 |
| 105 | Anterior | 0.367 ± 0.184 | 0.349 ± 0.356 | 0.115 ± 0.060 | 0.094 ± 0.112 | 0.182 ± 0.146 | 0.194 ± 0.184 | 0.024 ± 0.021 | 0.039 ± 0.041 |
| 105 | Posterior | 0.167 ± 0.111 | 0.000 ± 0.000 | 0.031 ± 0.029 | 0.000 ± 0.000 | 0.636 ± 0.303 | 0.765 ± 0.353 | 0.187 ± 0.091 | 0.251 ± 0.127 |
| 147 | Whole Bone | 0.572 ± 0.437 | 0.743 ± 0.539 | 0.247 ± 0.191 | 0.267 ± 0.225 | 0.932 ± 0.444 | 0.835 ± 0.218 | 0.392 ± 0.200 | 0.411 ± 0.143 |
| 147 | Anterior | 0.590 ± 0.461 | 0.683 ± 0.590 | 0.172 ± 0.138 | 0.205 ± 0.201 | 0.357 ± 0.282 | 0.297 ± 0.272 | 0.045 ± 0.044 | 0.050 ± 0.057 |
| 147 | Posterior | 0.092 ± 0.138 | 0.000 ± 0.000 | 0.016 ± 0.025 | 0.000 ± 0.000 | 0.940 ± 0.482 | 1.034 ± 0.188 | 0.240 ± 0.131 | 0.277 ± 0.068 |

**Supplemental Table 3. Trabecular and cortical bone parameters in 3-month-old osteoblast-specific *Rnf43* knockout mice.**

Micro-computed tomography (µCT) analysis of femoral trabecular and cortical bone from 3-month-old female and male control and osteoblast-specific *Rnf43* knockout mice. Sample sizes were as follows: females (control n = 9; knockout n = 11) and males (control n = 9; knockout n = 11). Data are presented as mean ± standard deviation (SD).

| 3 months-old *Ocn*-Cre-*Rnf43*$^{flox/flox}$ | Female | | Male | |
|---|---|---|---|---|
| Index | Control | *Rnf43* KO | Control | *Rnf43* KO |
| Trab.TV ($mm^3$) | 4.113 ± 0.901 | 4.142 ± 0.715 | 3.925 ± 0.487 | 3.943 ± 0.741 |
| Trab.BV ($mm^3$) | 0.464 ± 0.280 | 0.461 ± 0.342 | 0.475 ± 0.224 | 0.452 ± 0.373 |
| Trab.BV/TV (%) | 10.574 ± 4.249 | 10.347 ± 6.234 | 11.741 ± 4.898 | 10.383 ± 6.604 |
| Trab.N (1/mm) | 2.016 ± 0.728 | 1.958 ± 1.037 | 2.263 ± 0.769 | 1.970 ± 1.090 |
| Trab.Th (mm) | 0.052 ± 0.006 | 0.051 ± 0.005 | 0.050 ± 0.006 | 0.051 ± 0.005 |
| Trab.Sp (mm) | 0.236 ± 0.043 | 0.244 ± 0.050 | 0.219 ± 0.041 | 0.242 ± 0.049 |
| Trab.BMD (µgHA/$cm^3$) | 0.187 ± 0.042 | 0.181 ± 0.057 | 0.202 ± 0.042 | 0.183 ± 0.054 |
| Cort.TV ($mm^2$) | 0.959 ± 0.049 | 0.958 ± 0.053 | 1.152 ± 0.120 | 1.266 ± 0.132 |
| Cort.BV ($mm^2$) | 0.473 ± 0.029 | 0.463 ± 0.030 | 0.539 ± 0.044 | 0.590 ± 0.062 |
| Cort.BV/TV (%) | 49.279 ± 1.347 | 48.364 ± 1.365 | 46.894 ± 2.254 | 46.652 ± 2.691 |
| Cs.Th (mm) | 0.186 ± 0.008 | 0.181 ± 0.008 | 0.186 ± 0.009 | 0.193 ± 0.015 |
| Cort.TMD (µgHA/$cm^3$) | 1.110 ± 0.029 | 1.102 ± 0.032 | 1.099 ± 0.027 | 1.103 ± 0.029 |
| CAF (%) | 49.3 ± 1.3 | 48.4 ± 1.4 | 46.9 ± 2.3 | 46.7 ± 2.7 |
| T.Ar ($mm^2$) | 1.585 ± 0.081 | 1.583 ± 0.088 | 1.904 ± 0.198 | 2.092 ± 0.218 |
| B.Ar ($mm^2$) | 0.781 ± 0.048 | 0.766 ± 0.049 | 0.891 ± 0.073 | 0.975 ± 0.103 |
| B.Pm (mm) | 8.395 ± 0.243 | 8.441± 0.254 | 9.555 ± 0.632 | 10.110 ± 0.659 |

**Supplemental Table 4. Trabecular and cortical bone parameters in 6-month-old osteoblast-specific *Rnf43* knockout mice.**

Micro-computed tomography (µCT) analysis of femoral trabecular and cortical bone from 6-month-old female and male control and osteoblast-specific *Rnf43* knockout mice. Sample sizes were as follows: females (control n = 10; knockout n = 10) and males (control n = 8; knockout n = 8). Data are presented as mean ± standard deviation (SD).

| 6 months-old *Ocn*-Cre-*Rnf43*$^{flox/flox}$ | Female | | Male | |
|---|---|---|---|---|
| Index | Control | *Rnf43* KO | Control | *Rnf43* KO |
| Trab.TV ($mm^3$) | 3.171 ± 0.273 | 3.405 ± 0.334 | 4.251 ± 0.527 | 4.345 ± 0.589 |
| Trab.BV ($mm^3$) | 0.087 ± 0.058 | 0.134 ± 0.054 | 0.479 ± 0.161 | 0.511 ± 0.228 |
| Trab.BV/TV (%) | 2.714 ± 1.656 | 3.959 ± 1.577 | 11.274 ± 3.349 | 11.447 ± 3.890 |
| Trab.N (1/mm) | 0.523 ± 0.270 | 0.749 ± 0.274 | 1.952 ± 0.529 | 1.939 ± 0.566 |
| Trab.Th (mm) | 0.050 ± 0.008 | 0.053 ± 0.006 | 0.057 ± 0.007 | 0.059 ± 0.007 |
| Trab.Sp (mm) | 0.385 ± 0.035 | 0.359 ± 0.031 | 0.241 ± 0.029 | 0.251 ± 0.040 |
| Trab.BMD (µgHA/$cm^3$) | 0.117 ± 0.024 | 0.128 ± 0.024 | 0.182 ± 0.044 | 0.185 ± 0.043 |
| Cort.TV ($mm^2$) | 0.983 ± 0.079 | 1.018 ± 0.099 | 1.214 ± 0.134 | 1.229 ± 0.163 |
| Cort.BV ($mm^2$) | 0.513 ± 0.048 | 0.530 ± 0.064 | 0.573 ± 0.056 | 0.574 ± 0.070 |
| Cort.BV/TV (%) | 52.218 ± 2.722 | 52.158 ± 4.643 | 47.460 ± 4.738 | 46.867 ± 3.226 |
| Cs.Th (mm) | 0.203 ± 0.015 | 0.206 ± 0.022 | 0.195 ± 0.018 | 0.194 ± 0.015 |
| Cort.TMD (µgHA/$cm^3$) | 1.172 ± 0.026 | 1.171 ± 0.028 | 1.116 ± 0.033 | 1.117 ± 0.036 |
| CAF (%) | 52.2 ± 2.7 | 52.2 ± 4.6 | 47.5 ± 4.7 | 46.9 ± 3.2 |
| T.Ar ($mm^2$) | 1.625 ± 0.131 | 1.683 ± 0.164 | 2.006 ± 0.221 | 2.032 ± 0.269 |
| B.Ar ($mm^2$) | 0.848 ± 0.079 | 0.877 ± 0.106 | 0.947 ± 0.092 | 0.949 ± 0.116 |
| B.Pm (mm) | 8.343 ± 0.370 | 8.502 ± 0.489 | 9.748 ± 0.600 | 9.762 ± 0.790 |
